# *ggcoverage*: an R package to visualize and annotate genome coverage for various NGS data

**DOI:** 10.1101/2022.09.01.503744

**Authors:** Yabing Song, Jianbin Wang

## Abstract

**Summary:** Visualizing genome coverage is of vital importance to inspect and interpret various next-generation sequencing (NGS) data. However, existing tools that perform such visualizations remain inflexible, complicated, lack necessary annotations and the figures generated are not publication-quality. Here, we introduce *ggcoverage*, an R package with the grammar of graphics implemented in *ggplot2*, providing a flexible, programmable and user-friendly way to visualize genome coverage, multiple available annotations to adapt to different NGS data, and generating publication-quality plots.

**Availability and Implementation:** *ggcoverage* is an R package available via CRAN (https://cran.r-project.org/web/packages/ggcoverage/index.html) under MIT license. Support is available via GitHub (https://github.com/showteeth/ggcoverage/issues). Data used to generate figures are available on GitHub as example data.

## 1 Introduction

Visualizing genome coverage is of vital importance to inspect and interpret various next-generation sequencing (NGS) data. When analyzing whole-genome sequencing (WGS) data to get copy number variations (CNV), genome coverage plot can check for possible confounding factors, such as GC content bias, telomeres and centromeres proximity (Nguyen *et al*., 2006). When dealing with RNA-sequencing (RNA-seq) data, we can utilize genome coverage plot to inspect the gene or exon knockout efficiency, 5’ or 3’ bias of read coverage and visualize the read counts of differentially expressed genes, transcripts or exons (Conesa *et al*., 2016). In processing chromatin immunoprecipitation followed by sequencing (ChIP-seq) data, genome coverage plot can help to obtain and verify the peaks by comparing the signal of ChIP and input samples and visualize the relative distance between identified peaks and nearby genes (Park, 2009).

Despite its importance, existing tools to visualize genome coverage are often inflexible, complicated, lack necessary annotations and the figures generated are not publication-quality. For example, *UCSC genome browser* (Kent *et al*., 2002) and *IGV Browser* (Robinson *et al*., 2011) require file upload or data transmission, which usually takes time, and are not accessible programmatically. *Gviz* (Hahne and Ivanek, 2016) offers limited customization of plot aesthetics and themes. *GenVisR* (Skidmore *et al*., 2016) only offers gene and GC content annotations and the figure generated needs further modification. *karyoploteR* (Gel and Serra, 2017) focuses on visualizing chromosome ideogram, and is complicated to create genome coverage plot.

Here, we present *ggcoverage*, an R package providing a flexible, programmable and user-friendly way to visualize genome coverage, and multiple available annotations such as base and amino acid annotation, GC content annotation, gene structure annotation, transcript structure annotation, peak annotation and chromosome ideogram annotation to better inspect and interpret various NGS data. Furthermore, *ggcoverage* can generate publication-ready plots with the help of *ggplot2* (Wickham, 2016).

## 2 Implementation

The input file for *ggcoverage* can be in BAM, BigWig, BedGraph or tab-separated formats. For BAM files, *ggcoverage* can convert them to BigWig files with various normalization methods using *deeptools* (Ramírez *et al*., 2016). For tab-separated files, it should contain columns to specify chromosome, start, end, sample type and sample group. *ggcoverage* also requires additional files to annotate the genome coverage plot, such as FASTA file for GC content annotation, Gene transfer format (GTF) file for gene and transcript annotations and peak file for peak annotation.

To visualize and annotate genome coverage, *ggcoverage* introduces seven layers: `geom_coverage, geom_base, geom_gc, geom_gene, geom_transcript, geom_peak and geom_ideogram`. Besides these layers, *ggcoverage* also provides corresponding theme to beautify figures. `geom_coverage` will generate coverage plot of specified region for different samples, while the samples in the same group will have the same fill color or user-specified colors (Fig. 1A-D). When mark regions are provided, `geom_coverage` will also highlight these regions with specified colors and labels (Fig. 1C). `geom_base` is used to show base frequency and reference base for each locus, and it will also show amino acids of given region in *IGV* style (Fig. 1B). `geom_gc` will calculate and visualize GC content of every bin, and it will also add a line to indicate mean GC content or user-specified GC content (Fig. 1A). `geom_gene` will obtain all genes in given region and classify these genes to different groups to avoid overlap when plotting. In gene annotation, the arrow direction indicates the strand of genes, the height of different elements indicates different gene parts, the color of line indicates gene strand or user-specified group information (e.g., gene type) (Fig. 1A, C). `geom_transcript` is similar to `geom_gene`, but it shows all transcripts of a gene rather than the whole gene structure (Fig. 1D). `geom_peak` will show the peaks identified, so that the peaks and the nearby genes can be well visualized (Fig. 1C). `geom_ideogram` will show chromosome ideogram to illustrate the relative position of the displayed regions on the chromosome (Fig. 1A-D).

**Fig. 1.**
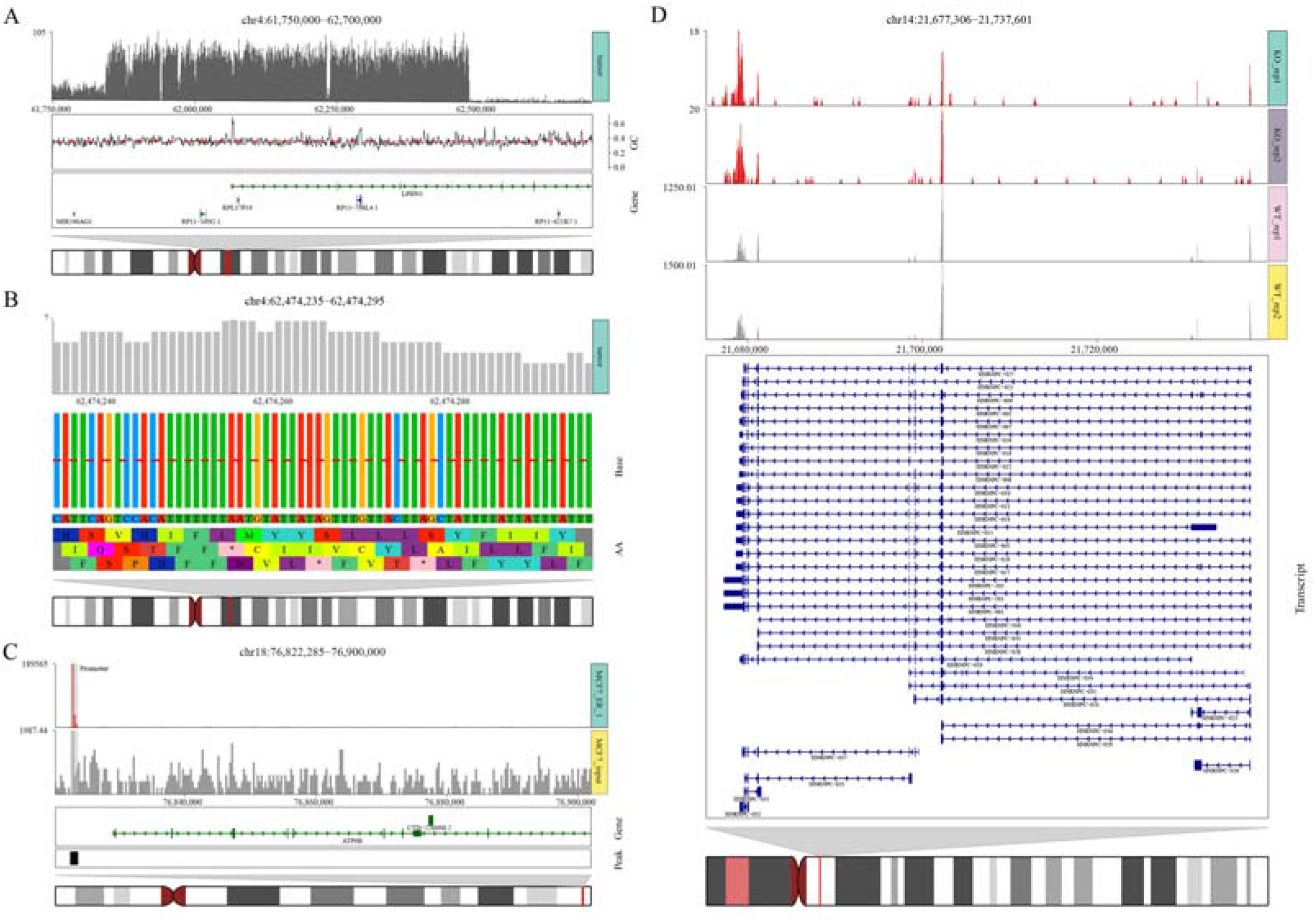
Visualizations of *ggcoverage* on selected NGS datasets. (**A**) *ggcoverage* on WGS data to visualize CNV. Genome coverage plot shows read counts of all bins. GC content annotation shows GC content of every bin (red line: mean GC content of the whole region). Gene annotation shows genes in given region (rightwards arrow with dark green: gene on plus strand, leftwards arrow with dark blue: gene on minus strand; element height: gene part (exon > UTR > intron)). Chromosome ideogram annotation shows the displayed region on the chromosome with red rectangle. (**B**) *ggcoverage* on WGS data to visualize single nucleotide variants (SNV). Genome coverage plot shows read counts of every locus. Base annotation shows base frequency (red line: 0.5) and reference base of every locus. Amino acid annotation shows corresponding amino acids with 0, 1, 2 offsets. (**C**) *ggcoverage* on ChIP-seq data. Different from (**A**), genome coverage plot discriminates sample groups (the first track in red is the ChIP sample and the last track in grey is the control sample), the light grey rectangle indicates the highlight region. Peak annotation shows all peaks identified. (**D**) *ggcoverage* on RNA-seq data with *HNRNPC* knockdown. Different from (**A**), we use transcript annotation instead of gene annotation to visualize gene’s all transcripts.

With the grammar of graphics implemented in *ggplot2*, users can superimpose the above layers by the `‘+’` operator. For example, `ggcoverage() + geom_gc() + geom_gene() + geom_ideogram()` will generate Fig. 1A. This elegant, concise and modular visualization system making it useful for genome coverage plot customization, curation and expansion.

## 3 Conclusion

We have developed *ggcoverage*, an R package dedicated to visualizing and annotating genome coverage. It allows users to visualize genome coverage with flexible input file formats, annotate the genome coverage with various annotations to meet the needs of different NGS technologies. And thanks to the multi-platform support of R, users do not need to transmit data. Finally, by using the grammar of graphics implemented in *ggplot2*, it is very convenient to generate high-quality and publication-ready plots.

